# Geometric Morphometrics of Pistillate Flowers in *Astrocaryum* sect. *Huicungo* (Arecaceae): Taxonomic Implications for the Peruvian Species

**DOI:** 10.64898/2026.11.04.764277

**Authors:** Valeria Meza, Gabriel Moreno, Marcelo Daza, Betty Millán

**Author notes:** Correspondence (B.M.). (G.M.); (V.M.); (M.D.).

## Abstract

The taxonomy of the Amazonian palm clade *Astrocaryum* sect. *Huicungo* remains controversial due to deep discordances between traditional classifications based on qualitative floral traits and recent phylogenomic evidence. To address this, landmark-based two-dimensional geometric morphometrics (4 landmarks, 16 semilandmarks) was applied to evaluate the symmetric component of shape variation in calyx and corolla structures across 549 pistillate flowers from the 11 Peruvian species. Although multivariate analyses (PCA, CVA) revealed continuous morphological variation and extensive phenotypic overlap, perianth geometry retained a significant diagnostic signal. Cross-validated classification rates demonstrated that the corolla exhibited higher discriminatory power than the calyx, while the combined perianth analysis (calyx + corolla) maximized taxonomic assignment accuracy. Furthermore, Mahalanobis distance-based clustering did not replicate traditional subsections: *A. chonta* and *A. ulei* exhibited divergent architectures, whereas *A. carnosum, A. faranae*, and *A. huicungo* shared profound geometric similarity. This study demonstrates that quantitative perianth morphometrics is a valuable tool for species delimitation in *Astrocaryum*, supporting the re-evaluation of traditional taxonomic boundaries through an integrative approach.

## 1. Introduction

The Arecaceae family constitutes an indispensable structural and functional component of tropical ecosystems across the Amazon basin and its peripheral regions, where it dominates various forest strata and provides vital resources for both wildlife and local communities **[1,2]**. Within this family, the genus *Astrocaryum* G. Mey. stands out for its high diversity, harboring approximately 40 species distributed primarily across South America. Section *Huicungo* (subgenus *Monogynanthus*) represents the most diverse lineage within the genus, comprising 15 species, 11 of which have their center of diversification in Peru **[3]**. These palms exhibit a remarkable variety of growth habits, ranging from acaulescent forms to large canopy-reaching arborescent palms, and occupy contrasting ecological niches, from seasonally flooded plains to terra firme forests and Andean foothills **[4,5]**.

Despite their ecological importance and hyperdominance, taxonomic delimitation within section *Huicungo* has been the subject of extensive and ongoing debate. The current infrageneric classification is primarily based on the qualitative evaluation of pistillate flower morphology, utilizing characters such as the length ratio between the calyx and corolla, the presence or absence of indumentum (prickles and bristles), and the architecture of the staminodial ring **[4]**. Based on these attributes, Kahn **[3]** subdivided the section into three subsections: *Huicungo, Sachacungo*, and *Murumuru*. However, current research has revealed strongly divergent hypotheses regarding the evolution of this group. On one hand, phylogenetic and phylogeographic studies have demonstrated a spatial structuring into two major geologically divided clades **[6,7]**. On the other hand, recent phylogenomic analyses based on plastid genomes have refuted the monophyly of the traditional subsections, suggesting that classifications based on qualitative floral morphology might be artificial and that taxonomic boundaries are highly diffuse **[8]**.

Given the discordance between phenotype and genotype, combined with the proven inadequacy of classical linear morphometrics to resolve species complexes with high intraspecific variability, geometric morphometrics has emerged as a crucial complementary quantitative approach **[9,10]**. This methodology, based on configurations of Cartesian coordinates (landmarks and semilandmarks), enables the isolation of pure geometric information by mathematically removing the effects of size, position, and rotation **[11]**. In botanical systematics, geometric morphometrics has proven highly effective for delimiting cryptic taxa from herbarium specimens, assessing evolutionary allometry, and quantifying the integration and shape of floral structures, thereby overcoming the inherent subjectivity of historical qualitative descriptions **[12,13]**.

Consequently, the main objective of this study is to quantitatively evaluate the morphological variability of the calyx and corolla shape of pistillate flowers in the Peruvian species of *Astrocaryum* sect. *Huicungo* using two-dimensional geometric morphometrics. This research seeks to determine whether perianth geometry retains a significant taxonomic signal that improves species delimitation, thus testing the viability of the classical classification system. The principal conclusions of this work reveal that, despite the presence of extensive phenotypic overlap and high floral developmental instability (fluctuating asymmetry), the combined geometry of the calyx and corolla retains a significant diagnostic signal. Furthermore, the recovered phenetic affinities do not replicate traditional subsections, providing quantitative evidence that aligns with recent phylogenomic and biogeographical hypotheses for the taxon.

## 2. Results

### 2.1. Shape Variation of the Calyx and Corolla Among Species

The Procrustes analysis of variance to evaluate bilateral symmetry (Table 1) revealed that the highest proportion of morphological variation in the perianth corresponds to the symmetric component (SC), which accounted for 80.07% of the total variance in the calyx and 68.08% in the corolla. The predominance and high statistical significance of this component (*p* < 0.001) confirm it as the primary source of geometric variation, thereby justifying the exclusive use of the symmetric component for all subsequent analyses.

**Table 1.**
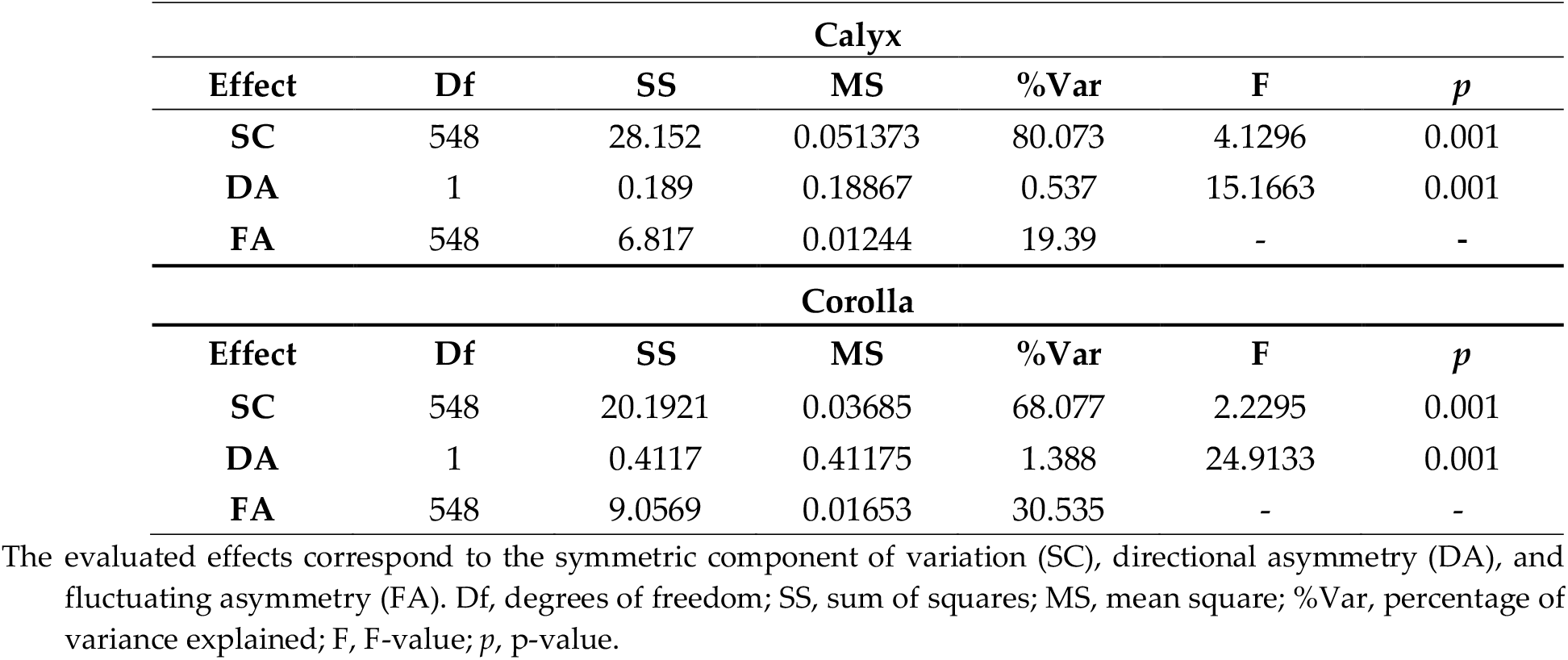
Results of the bilateral symmetry analysis for the calyx and corolla outline.

| **Calyx** |  |  |  |  |  |  |
|---|---|---|---|---|---|---|
| **Effect** | **Df** | **SS** | **MS** | **%Var** | **F** | ***p*** |
| **SC** | 548 | 28.152 | 0.051373 | 80.073 | 4.1296 | 0.001 |
| **DA** | 1 | 0.189 | 0.18867 | 0.537 | 15.1663 | 0.001 |
| **FA** | 548 | 6.817 | 0.01244 | 19.39 | - | **-** |
| **Corolla** |  |  |  |  |  |  |
| **Effect** | **Df** | **SS** | **MS** | **%Var** | **F** | ***p*** |
| **SC** | 548 | 20.1921 | 0.03685 | 68.077 | 2.2295 | 0.001 |
| **DA** | 1 | 0.4117 | 0.41175 | 1.388 | 24.9133 | 0.001 |
| **FA** | 548 | 9.0569 | 0.01653 | 30.535 | - | - |

The evaluated effects correspond to the symmetric component of variation (SC), directional asymmetry (DA), and fluctuating asymmetry (FA). Df, degrees of freedom; SS, sum of squares; MS, mean square; %Var, percentage of variance explained; F, F-value; *p*, p-value.

Conversely, fluctuating asymmetry (FA) constituted the second most important factor of variation, capturing 19.39% of the variance in the calyx and 30.53% in the corolla. This proportion evidences a considerable degree of intra-individual developmental instability in response to random perturbations. Finally, although directional asymmetry (DA) accounted for a marginal fraction of the total variance (0.53% for the calyx and 1.38% for the corolla), its effect was statistically significant (*p* < 0.001), indicating the existence of a subtle yet consistent systematic differentiation between the sides of both structures at the population level.

To further quantify the relative contribution of taxonomic, intra-specific, and architectural factors to floral shape variation, a hierarchical Procrustes ANOVA was performed (Table 2). The analysis revealed that species identity constituted the primary source of geometric variation, explaining 46.90% and 44.00% of the variance in the calyx and corolla, respectively (*p* = 0.001). This confirms that the perianth retained a robust diagnostic taxonomic signal. Notably, intra-specific variability (individuals nested within species) also accounted for a substantial proportion of the variance (31.00% in the calyx and 27.30% in the corolla), quantitatively corroborating the high phenotypic plasticity and developmental instability of these structures. Conversely, the architectural effect—defined by the position of the flower on the rachilla (apical, middle, or basal)—exerted a statistically significant but biologically marginal influence, explaining only 0.60% and 2.80% of the shape variance in the calyx and corolla, respectively.

**Table 2.**
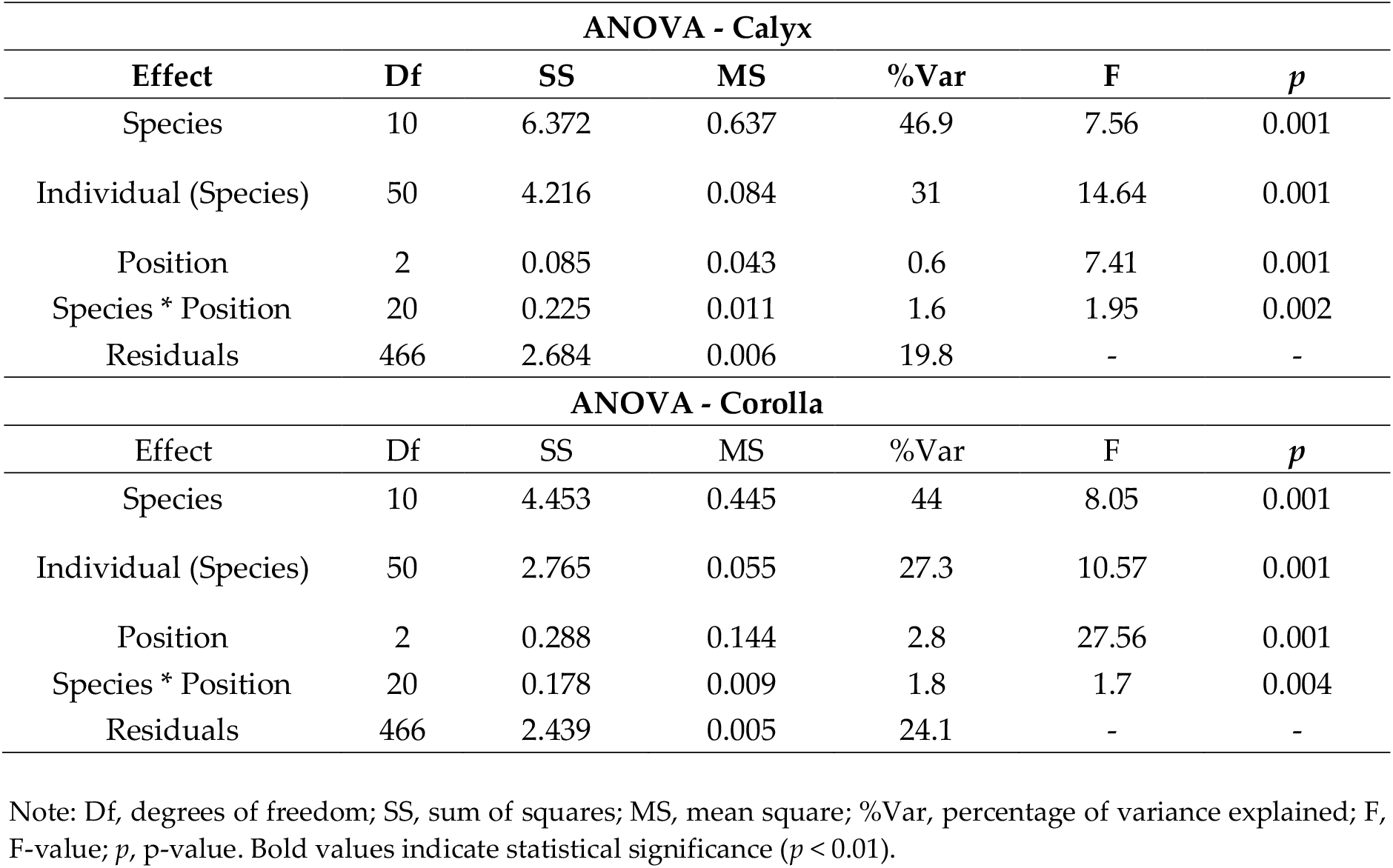
Hierarchical Procrustes ANOVA evaluating the effects of species, floral position, and individual variation on the symmetric shape component of the calyx and corolla.

The Principal Component Analysis (PCA) of the calyx revealed that the first two components (PC1 and PC2) accounted for 94.41% of the total variation (Figure 1a). Despite a pronounced general overlap, *Astrocaryum ulei* and *A. chonta* were distinguished by a clear trend toward positive values on both axes. In the corolla, PC1 and PC2 explained 89.21% of the morphological variance (Figure 1b), showing a high degree of morphospace overlap. The combined analysis (calyx + corolla) (Figure 1c) demonstrated that the first two components captured 85.65% of the geometric variation, displaying an ordination pattern consistent with that of the calyx.

**Figure 1.**
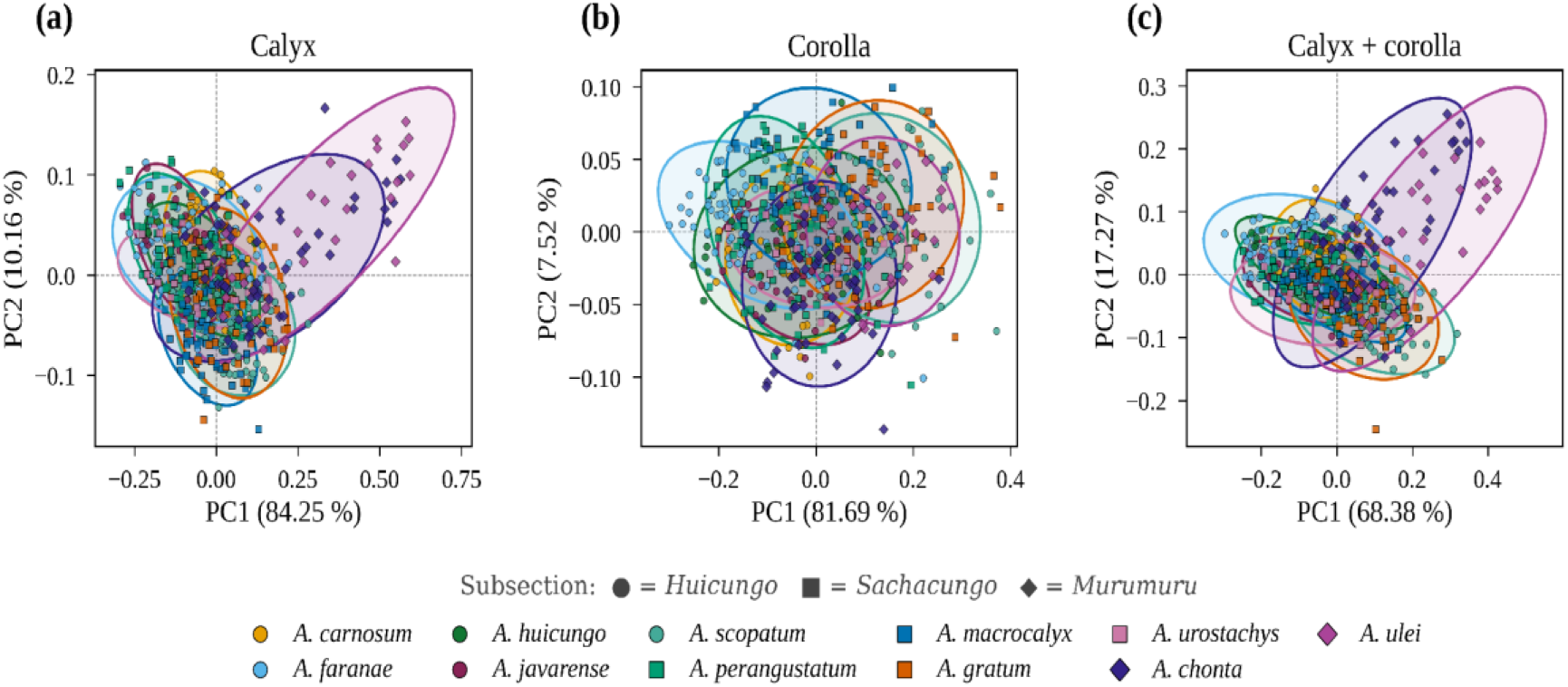
PCA plots utilizing the symmetric component of shape, considering all species of the *Huicungo* Section. Each point represents an individual pistillate flower, and the shaded confidence ellipses are drawn at a 95% probability level. **(a)** Calyx; **(b)** Corolla; **(c)** Calyx + corolla.

Upon evaluating the symmetric shape changes (Figure 2), variation along PC1 in the calyx represented a gradient from elongated, narrow shapes (associated with negative values) to broad, short structures (positive values). The variation explained by PC2 was concentrated at the base and distal margin; specifically, positive values denoted a widening at the base accompanied by distal narrowing (Figure 2a).

**Figure 2.**
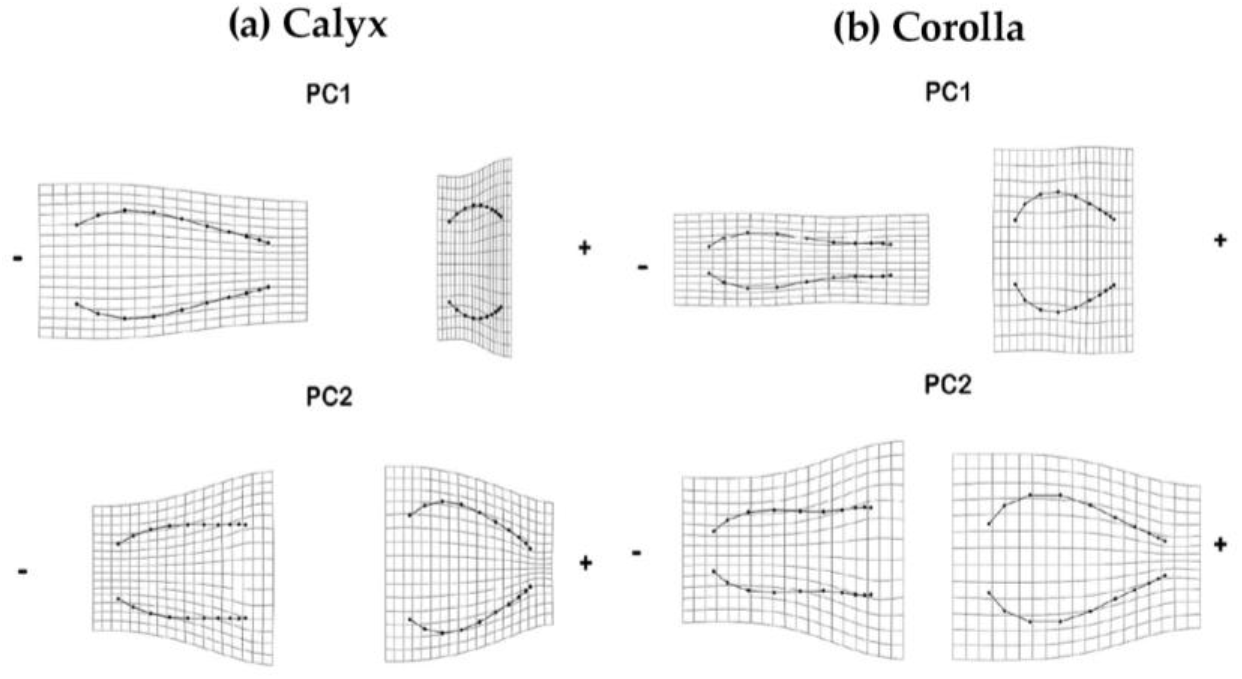
Symmetric shape changes corresponding to the extreme values (negative and positive) of PC1 and PC2. (a) Calyx; (b) Corolla.

Congruently, the variation patterns in the corolla replicated the trends observed in the calyx. PC1 described the same longitudinal and transverse morphological transition (from elongated/narrow to broad/short). Finally, PC2 modeled the transverse variation at the corolla extremities: the negative pole displayed a conspicuously narrow base with a broad distal margin, whereas the positive pole exhibited the inverse conformation (broad base and narrow distal margin) (Figure 2b).

### 2.2. Mean Shapes of the Calyx and Corolla

The mean shapes of the calyx and corolla, estimated via Procrustes analysis, demonstrated high overall similarity among the 11 studied species, exhibiting only minor deviations from the consensus shapes. Most calyx conformations represented variants of a tubular shape, whereas the corollas tended to be narrower, ranging from tubular to urceolate or obovoid forms (Figure 3).

**Figure 3.**
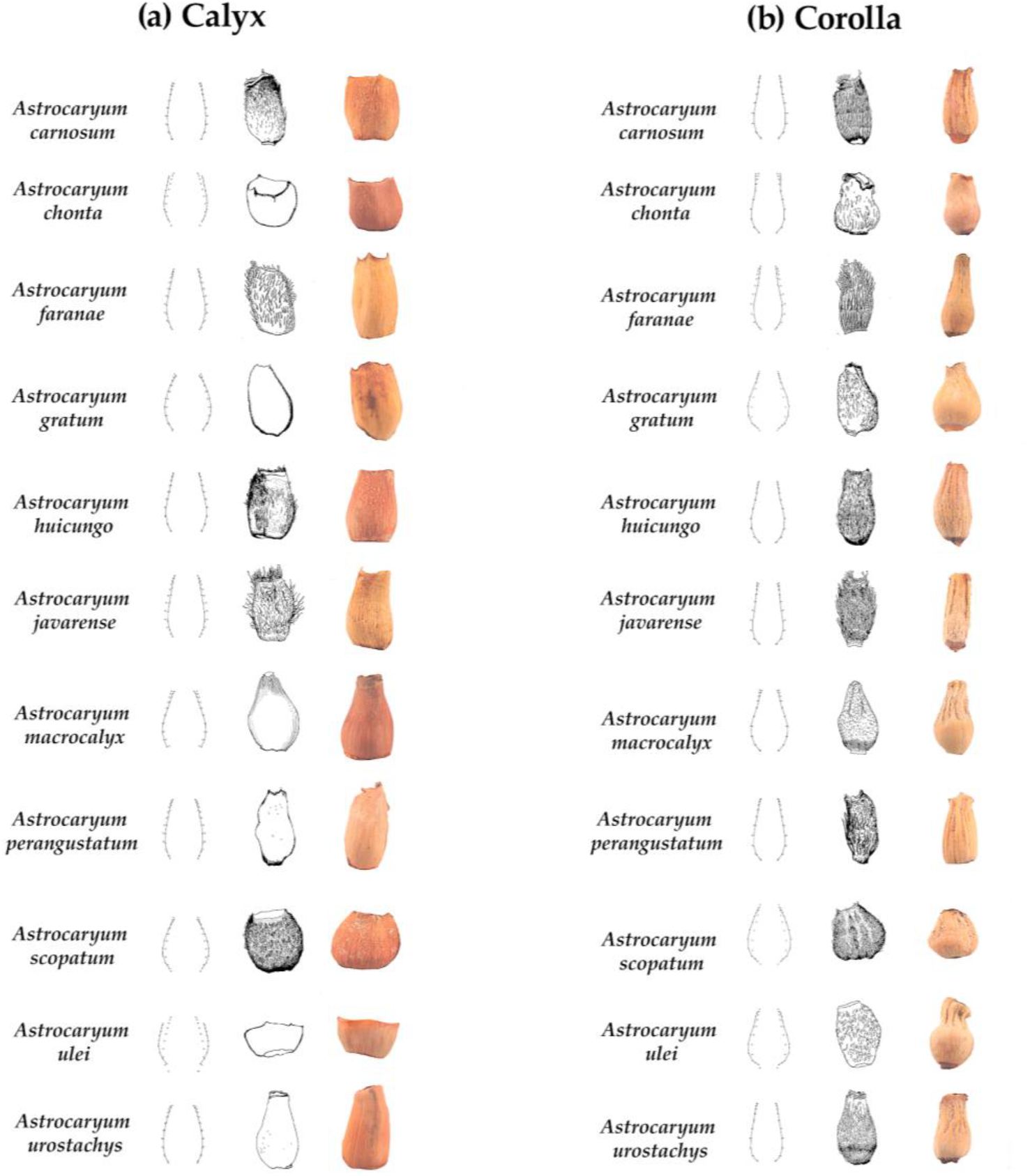
Estimated mean shapes for the floral parts of the *Huicungo* Section species. From left to right: the mean shape calculated using the symmetric component for each species and its deviation from the consensus; graphic representation according to the descriptions by Kahn and Millán [14] and Kahn and Ferreira [15]; and a photograph of a representative calyx for each species (Meza, V. and Moreno, G.). (a) Calyx; (b) Corolla.

Visualization of the mean calyx shape (Figure 3a) in *Astrocaryum perangustatum* stood out for exhibiting a clearly tubular morphology with a slender, cylindrical profile, which coincided with the negative extreme of PC1. Similarly, *A. javarense* and *A. faranae* presented markedly narrow, elongated calyces; however, *A. javarense* did not exhibit a perfectly oblong shape. In contrast, *A. ulei* and *A. chonta* were characterized by shorter calyces with a broad distal margin. Although both presented an affinity toward cupuliform shapes, *A. chonta* exhibited a slightly more elongated conformation than *A. ulei. A. carnosum* stood out due to a slightly widened distal margin. Toward the positive extreme of PC1, *A. gratum* displayed a characteristically urceolate morphology—notably ventricose at the base and slightly narrowed toward the apex. *A. scopatum* shared this tendency toward a broad base and a narrow orifice, but with a markedly ovoid conformation. Finally, *A. macrocalyx* showed a suburceolate tendency, presenting an elongated structure that was more ventricose in the medial region than the tubular taxa.

In the corolla (Figure 3b), the aspect ratio and lateral curvature allowed for the differentiation of consistent morphological groups. *Astrocaryum carnosum* and *A. huicungo* presented highly similar conformations with minimal deviations. As observed in the calyx, *A. perangustatum* defined the variation extreme toward slender shapes, characterized by an elongated, tubular outline lacking basal expansion—a trend also exhibited by *A. faranae*. The species *A. ulei, A. gratum*, and *A. scopatum* were characterized by notably broad bases and narrow orifices. Specifically, *A. scopatum* retained the ovoid shape observed in its calyx, while *A. gratum* exhibited a pyriform conformation with an evident basal expansion that gradually narrowed toward the apex. *A. urostachys* presented a globose to ovoid morphology, comparatively broader and with an abruptly truncated apex. Regarding the distal margin, *A. chonta* maintained a basal urceolate shape but featured the broadest margin of the group. Conversely, *A. macrocalyx* presented the narrowest distal margin, standing out for its regular, symmetrical outline and a slight subapical constriction.

### 2.3. Species Delimitation and Discriminatory Power

The Canonical Variate Analysis (CVA) corroborated the extensive phenotypic overlap among the studied taxa. When evaluating the structures in isolation, the CV1 and CV2 axes accounted for 66.93% of the variance for the calyx and 72.18% for the corolla. Conversely, the combined analysis (calyx + corolla) captured 61.01% of the geometric variation (Figure 4).

**Figure 4.**
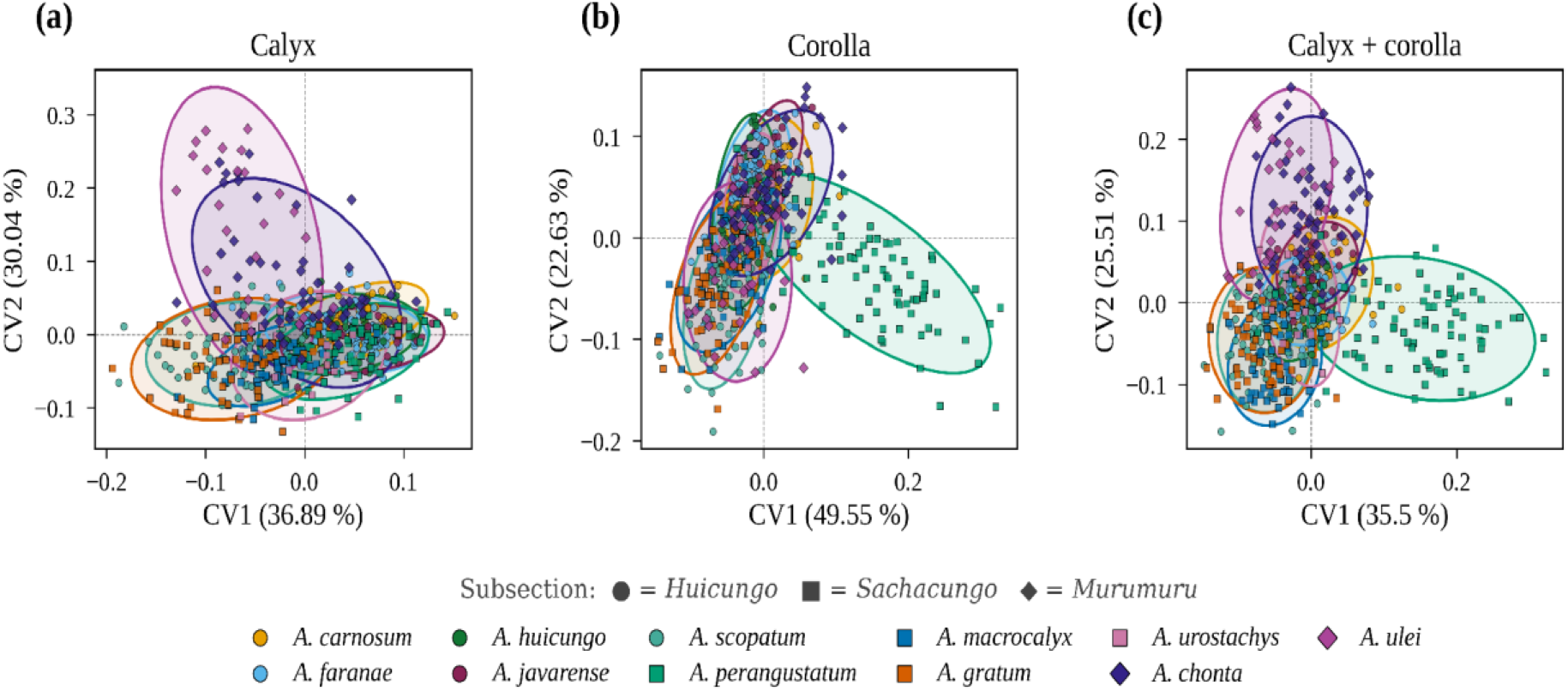
CVA plots utilizing the symmetric component of floral part shape for the species of the *Huicungo* Section. Each point represents an individual pistillate flower, and the shaded confidence ellipses are drawn at a 95% probability level. (a) Calyx; (b) Corolla; (c) Calyx + corolla.

In the specific analysis of the calyx, this high morphospace superimposition precluded an absolute distinction among taxa. Nonetheless, CV2 partially segregated *Astrocaryum chonta* and *A. ulei* from the remaining species.

Regarding the corolla, despite the high degree of overlap that hindered the unequivocal recognition of the groups, CV1 contributed to the partial separation of *Astrocaryum perangustatum* from the other species, while CV2 differentiated *A. chonta* from *A. ulei*.

Finally, the combined analysis (calyx + corolla) also exhibited a high degree of overlap; consistent with the corolla-only analysis, CV1 contributed to the partial segregation of *Astrocaryum perangustatum* from the remaining taxa.

The cross-validated classification test derived from the Canonical Variate Analysis (CVA) yielded a correct classification rate (CCR) of 53.92% for the corolla and 44.26% for the calyx. The combined perianth dataset (calyx + corolla) resulted in a CCR of 64.85%. Based on the pairwise Mahalanobis distances evaluated with a Bonferroni correction (Table 3), the calyx recorded significant morphological differentiation (*p* < 0.05) across most species comparisons. However, no significant differences (*p* > 0.05) were detected between *A. perangustatum* and *A. urostachys*, nor among *Astrocaryum carnosum, A. faranae, A. huicungo*, and *A. javarense*.

**Table 3.**
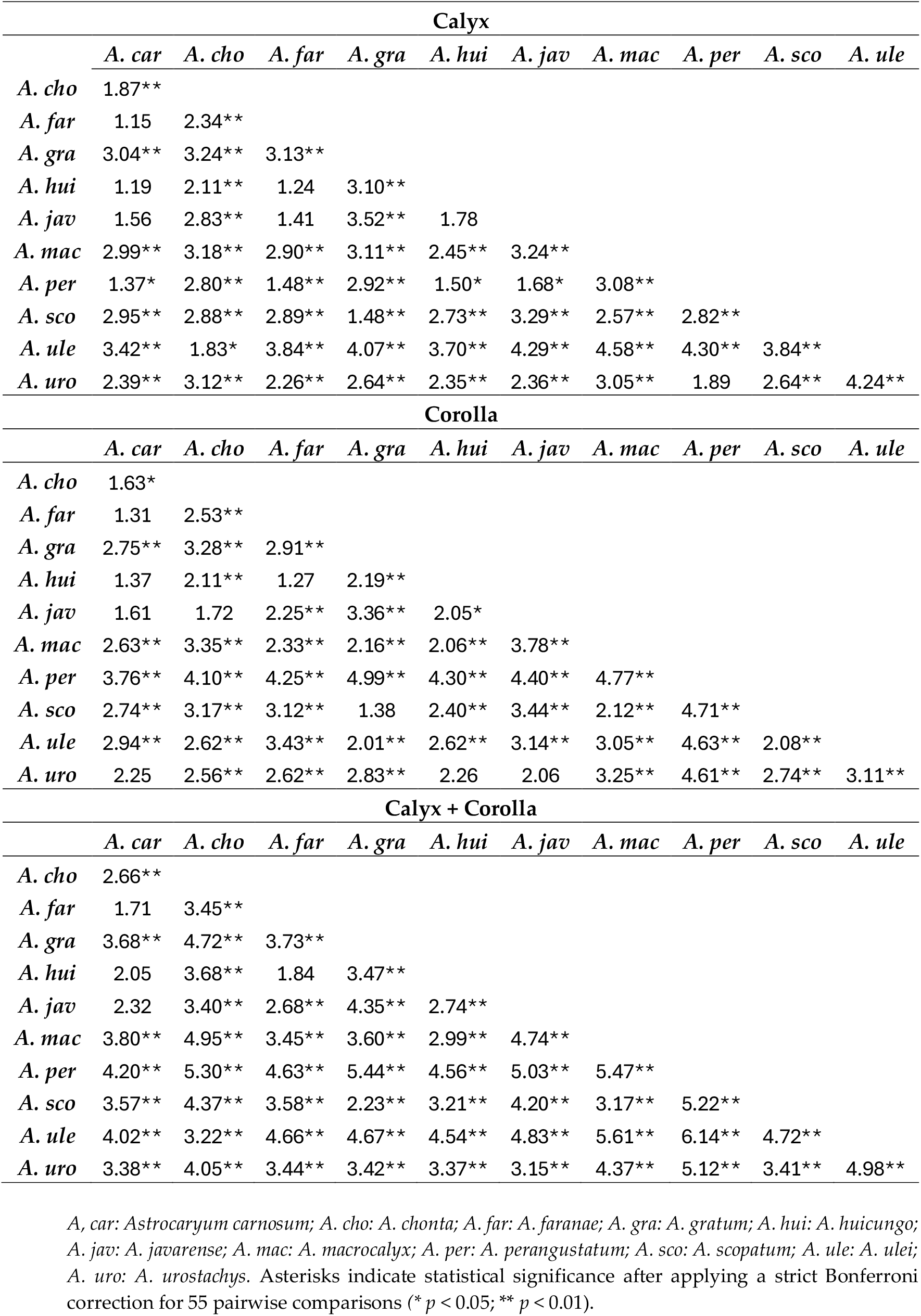
Pairwise Mahalanobis distances derived from the Canonical Variate Analysis (CVA) of the symmetric shape component for the calyx, corolla, and combined perianth datasets.

| **Calyx** |  |  |  |  |  |  |  |  |  |  |
|---|---|---|---|---|---|---|---|---|---|---|
|  | *A. car* | *A. cho* | *A. far* | *A. gra* | *A. hui* | *A. jav* | *A. mac* | *A. per* | *A. sco* | *A. ule* |
| ***A. cho*** | 1.87** |  |  |  |  |  |  |  |  |  |
| ***A. far*** | 1.15 | 2.34** |  |  |  |  |  |  |  |  |
| ***A. gra*** | 3.04** | 3.24** | 3.13** |  |  |  |  |  |  |  |
| ***A. hui*** | 1.19 | 2.11** | 1.24 | 3.10** |  |  |  |  |  |  |
| ***A. jav*** | 1.56 | 2.83** | 1.41 | 3.52** | 1.78 |  |  |  |  |  |
| ***A. mac*** | 2.99** | 3.18** | 2.90** | 3.11** | 2.45** | 3.24** |  |  |  |  |
| ***A. per*** | 1.37* | 2.80** | 1.48** | 2.92** | 1.50* | 1.68* | 3.08** |  |  |  |
| ***A. sco*** | 2.95** | 2.88** | 2.89** | 1.48** | 2.73** | 3.29** | 2.57** | 2.82** |  |  |
| ***A. ule*** | 3.42** | 1.83* | 3.84** | 4.07** | 3.70** | 4.29** | 4.58** | 4.30** | 3.84** |  |
| ***A. uro*** | 2.39** | 3.12** | 2.26** | 2.64** | 2.35** | 2.36** | 3.05** | 1.89 | 2.64** | 4.24** |
| **Corolla** |  |  |  |  |  |  |  |  |  |  |
|  | *A. car* | *A. cho* | *A. far* | *A. gra* | *A. hui* | *A. jav* | *A. mac* | *A. per* | *A. sco* | *A. ule* |
| ***A. cho*** | 1.63* |  |  |  |  |  |  |  |  |  |
| ***A. far*** | 1.31 | 2.53** |  |  |  |  |  |  |  |  |
| ***A. gra*** | 2.75** | 3.28** | 2.91** |  |  |  |  |  |  |  |
| ***A. hui*** | 1.37 | 2.11** | 1.27 | 2.19** |  |  |  |  |  |  |
| ***A. jav*** | 1.61 | 1.72 | 2.25** | 3.36** | 2.05* |  |  |  |  |  |
| ***A. mac*** | 2.63** | 3.35** | 2.33** | 2.16** | 2.06** | 3.78** |  |  |  |  |
| ***A. per*** | 3.76** | 4.10** | 4.25** | 4.99** | 4.30** | 4.40** | 4.77** |  |  |  |
| ***A. sco*** | 2.74** | 3.17** | 3.12** | 1.38 | 2.40** | 3.44** | 2.12** | 4.71** |  |  |
| ***A. ule*** | 2.94** | 2.62** | 3.43** | 2.01** | 2.62** | 3.14** | 3.05** | 4.63** | 2.08** |  |
| ***A. uro*** | 2.25 | 2.56** | 2.62** | 2.83** | 2.26 | 2.06 | 3.25** | 4.61** | 2.74** | 3.11** |
| **Calyx + Corolla** |  |  |  |  |  |  |  |  |  |  |
|  | *A. car* | *A. cho* | *A. far* | *A. gra* | *A. hui* | *A. jav* | *A. mac* | *A. per* | *A. sco* | *A. ule* |
| ***A. cho*** | 2.66** |  |  |  |  |  |  |  |  |  |
| ***A. far*** | 1.71 | 3.45** |  |  |  |  |  |  |  |  |
| ***A. gra*** | 3.68** | 4.72** | 3.73** |  |  |  |  |  |  |  |
| ***A. hui*** | 2.05 | 3.68** | 1.84 | 3.47** |  |  |  |  |  |  |
| ***A. jav*** | 2.32 | 3.40** | 2.68** | 4.35** | 2.74** |  |  |  |  |  |
| ***A. mac*** | 3.80** | 4.95** | 3.45** | 3.60** | 2.99** | 4.74** |  |  |  |  |
| ***A. per*** | 4.20** | 5.30** | 4.63** | 5.44** | 4.56** | 5.03** | 5.47** |  |  |  |
| ***A. sco*** | 3.57** | 4.37** | 3.58** | 2.23** | 3.21** | 4.20** | 3.17** | 5.22** |  |  |
| ***A. ule*** | 4.02** | 3.22** | 4.66** | 4.67** | 4.54** | 4.83** | 5.61** | 6.14** | 4.72** |  |
| ***A. uro*** | 3.38** | 4.05** | 3.44** | 3.42** | 3.37** | 3.15** | 4.37** | 5.12** | 3.41** | 4.98** |

*A, car: Astrocaryum carnosum; A. cho: A. chonta; A. far: A. faranae; A. gra: A. gratum; A. hui: A. huicungo; A. jav: A. javarense; A. mac: A. macrocalyx; A. per: A. perangustatum; A. sco: A. scopatum; A. ule: A. ulei; A. uro: A. urostachys*. Asterisks indicate statistical significance after applying a strict Bonferroni correction for 55 pairwise comparisons (\* *p* < 0.05; \*\* *p* < 0.01).

Regarding the corolla, 9 species pairs lacked significant morphological differentiation (*p* > 0.05), including the pairwise comparisons among *A. carnosum, A. faranae*, and *A. huicungo*. Finally, the combined analysis (calyx + corolla) yielded significant Mahalanobis distances for all interspecific comparisons, with the exception of the pairwise comparisons among *Astrocaryum carnosum, A. faranae*, and *A. huicungo*, which maintained non-significant distances (*p* > 0.05) among themselves. Notably, the combined perianth geometry successfully discriminated *A. javarense* from this group, evidencing highly significant morphometric distances (*p* < 0.01) against *A. faranae* and *A. huicungo*

The cluster analysis based on Mahalanobis distances for the calyx (Figure 5a) revealed that *Astrocaryum ulei* constitutes the most morphologically divergent taxon, positioning itself isolated from the remaining species. Within the main cluster, two primary groups were distinguished: the first comprised *A. scopatum, A. gratum*, and *A. macrocalyx*, united by geometric proximity; the second was characterized by high similarity among *A. carnosum, A. huicungo*, and *A. faranae*, forming a subgroup to which *A. perangustatum, A. javarense, A. urostachys*, and *A. chonta* adhered successively.

**Figure 5.**
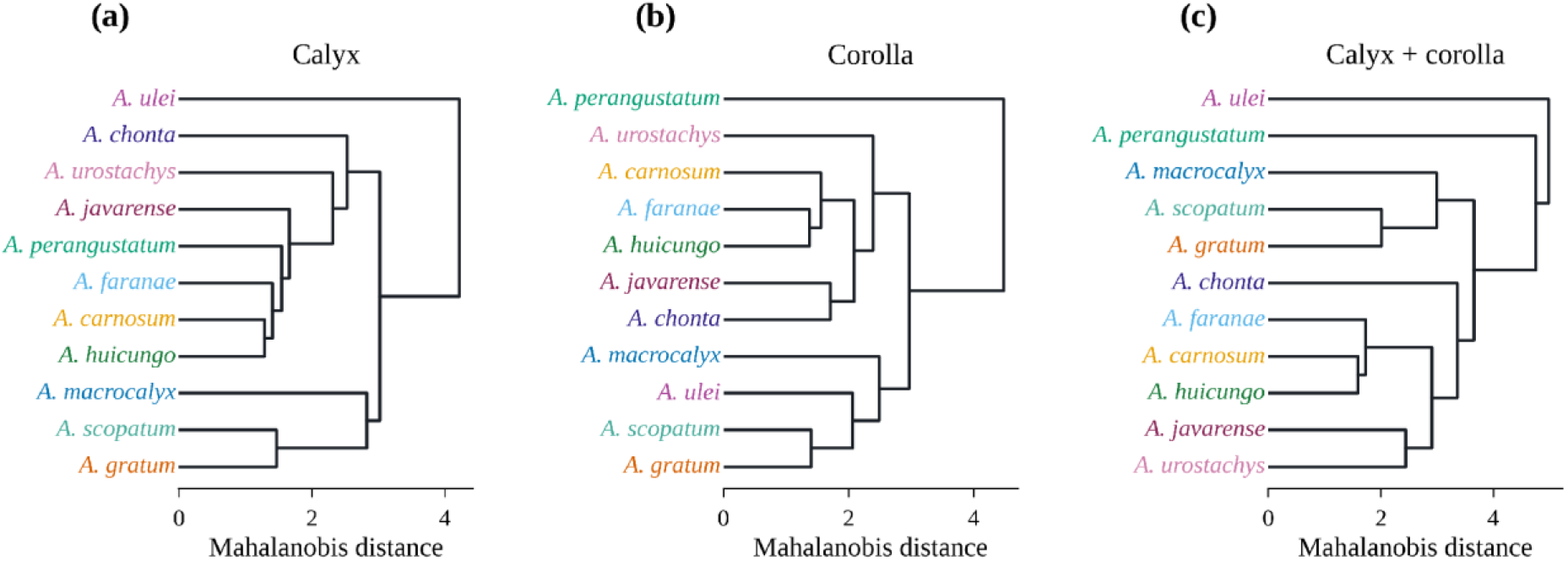
Similarity dendrogram obtained from Mahalanobis distances. (a) Calyx; (b) Corolla; (c) calyx + corolla.

Regarding the corolla conformation (Figure 5b), the similarity pattern exhibited notable variations. In this case, *Astrocaryum perangustatum* was recovered as the most divergent species. The group formed by *A. scopatum* and *A. gratum* maintained its proximity, but on this occasion integrated *A. ulei* and *A. macrocalyx*. Conversely, the close morphological affinity among *A. carnosum, A. faranae*, and *A. huicungo* remained consistent, forming a cluster associated with *A. urostachys*, while *A. javarense* and *A. chonta* formed a subgroup close to the latter block.

Finally, the dendrogram calculated from the concatenated dataset (calyx + corolla) synthesized the geometric trends of both structures (Figure 5c). *Astrocaryum ulei* and *A. perangustatum* consolidated as the most divergent species, branching externally to the rest of the group. The cluster integrated by *A. scopatum, A. gratum*, and *A. macrocalyx* was recovered, consistent with the observation for the isolated calyx. Likewise, the high phenotypic affinity among *A. carnosum, A. huicungo*, and *A. faranae* remained invariable, constituting the core of a broader block that sequentially grouped *A. javarense, A. urostachys*, and *A. chonta*.

## 3. Discussion

### 3.1. Symmetry and Developmental Instability in Floral Morphology

A fundamental biological finding derived from the Procrustes superimposition was the quantification of object bilateral symmetry. Although the symmetric component (SC) accounted for the largest proportion of the geometric variance, a high percentage of fluctuating asymmetry (FA) was detected, capturing 19.39% of the variation in the calyx and 30.53% in the corolla. In geometric morphometrics, fluctuating asymmetry represents small, random deviations from perfect symmetry and is considered a direct indicator of instability or developmental noise during ontogeny **[16]**. This level of morphological instability is not unusual in dense inflorescences and has been corroborated in other complex botanical families, such as Ranunculaceae **[17]**, as well as in the foliar variation of the genus *Alnus*, where fluctuating asymmetry reflects high phenotypic plasticity in response to environmental stressors **[18]**.

Because fluctuating asymmetry is not genetically determined, its high prevalence indicates a strong influence of microenvironmental factors. Traditional models often attribute this variation to the position of the flower on the rachilla and local resource allocation **[19,20]**. However, our hierarchical ANOVA demonstrated that flower position had a marginal impact, explaining only 0.60% to 2.80% of the shape variance. Instead, this developmental instability is likely driven by physical crowding as the flowers grow tightly packed on the rachilla, forcing them into constrained developmental modules **[21]**.

In addition to these ontogenetic pressures, the perianth faces intense ecological interactions. The chewing activity of pollinating beetles, which is extensively documented in the genus **[22]**, imposes severe mechanical stress that could exacerbate the observed asymmetry and plasticity in mature floral stages. Collectively, the convergence of these intrinsic constraints and extrinsic pressures confirms that the final floral shape in this group is a highly plastic trait, thereby contributing to the extensive interspecific overlap documented in the multivariate analyses (PCA and CVA).

### 3.2. Geometric Variation and Phenotypic Overlap

Geometric morphometric analysis revealed a continuous morphological variation in the perianth of *Astrocaryum* sect. *Huicungo* pistillate flowers, characterized by an extensive phenotypic overlap within the morphospace. This high intra- and interspecific variability challenges the discrete categorization of floral shapes traditionally employed in the taxonomy of the group **[4]**. The use of broad qualitative terminology (e.g., tubular, urceolate, dolioliform) may oversimplify floral geometry and underestimate the true variation present in complex taxa **[23]**. In this context, geometric morphometrics has proven to be a superior tool for isolating biological shape and detecting subtle phenotypic variations that linear measurements fail to capture **[9,24,25]**.

Our mean shape estimations demonstrate that, while identifiable extreme phenotypes exist—such as the distinctive urceolate calyx of *A. gratum* or the tubular conformation of *A. perangustatum*—most species exhibit redundant floral conformations. This aligns with biometric observations of extensive inter- and intrapopulation variation previously reported for the genus **[26]**.

### 3.3. Taxonomic Delimitation and Phylogenetic Discordance

Despite the continuous variation and visual overlap observed in the Principal Component Analyses (PCA), statistical tests confirmed that perianth geometry retains a significant diagnostic signal. Specifically, the cross-validated classification rates demonstrated that combining the calyx and corolla datasets substantially enhances taxonomic discrimination, achieving the highest correct classification rate (64.85%). This improvement achieved by concatenating multiple morphological structures aligns with documentation in other complex botanical groups, where multi-structure integration more faithfully reflects evolutionary lineages and maximizes phenotypic resolution **[27,28]**.

However, the interpretation of these statistical differences requires analytical caution. On one hand, utilizing individual flowers as sampling units can introduce pseudoreplication effects that may artificially inflate statistical significance **[29]**. On the other hand, the high dimensionality inherent to geometric morphometric coordinate datasets can generate apparent mathematical distances even among highly similar morphological distributions **[30,31]**. Therefore, while the pairwise Mahalanobis distances support the separation of most species, these metrics must be understood as mathematical indicators of shape divergence rather than absolute biological barriers.

Beyond these analytical considerations, phenetic similarity relationships derived from the UPGMA clustering revealed a clear incongruence with the taxonomic subsections proposed by Kahn **[3]**. A representative case is the high phenetic proximity observed in the UPGMA clustering among *Astrocaryum carnosum, A. faranae*, and *A. huicungo*. However, the interpretation of these phenotypic affinities must decouple graphic clustering from absolute statistical non-significance; indeed, the concatenated pairwise Mahalanobis distances revealed that the combined perianth successfully discriminates *A. carnosum* from its counterparts, whereas non-significant differences persist strictly within the isolated calyx analysis. This shared geometric pattern in the calyx structure points toward limited morphological differentiation or a recent divergence event rather than evolutionary convergence, a scenario highly congruent with recent plastid genome phylogenomics that failed to recover the monophyly of traditional subsections **[7,8]**. Ultimately, this complex morphological continuum among geographically contiguous western Amazonian taxa **[32]** likely reflects a process of recent speciation where ancestral genetic traits are still shared, driven by Late Neogene geoclimatic transitions **[33]**.

## 4. Materials and Methods

### 4.1. Plant material

The botanical material for this study was obtained from herbarium specimens deposited at the San Marcos Herbarium (USM), representing the 11 species of the genus *Astrocaryum* sect. *Huicungo* found in Peru (*Astrocaryum carnosum, A. chonta, A. faranae, A. gratum, A. huicungo, A. javarense, A. macrocalyx, A. perangustatum, A. scopatum, A. ulei*, and *A. urostachys*) (Figure 6).

**Figure 6.**
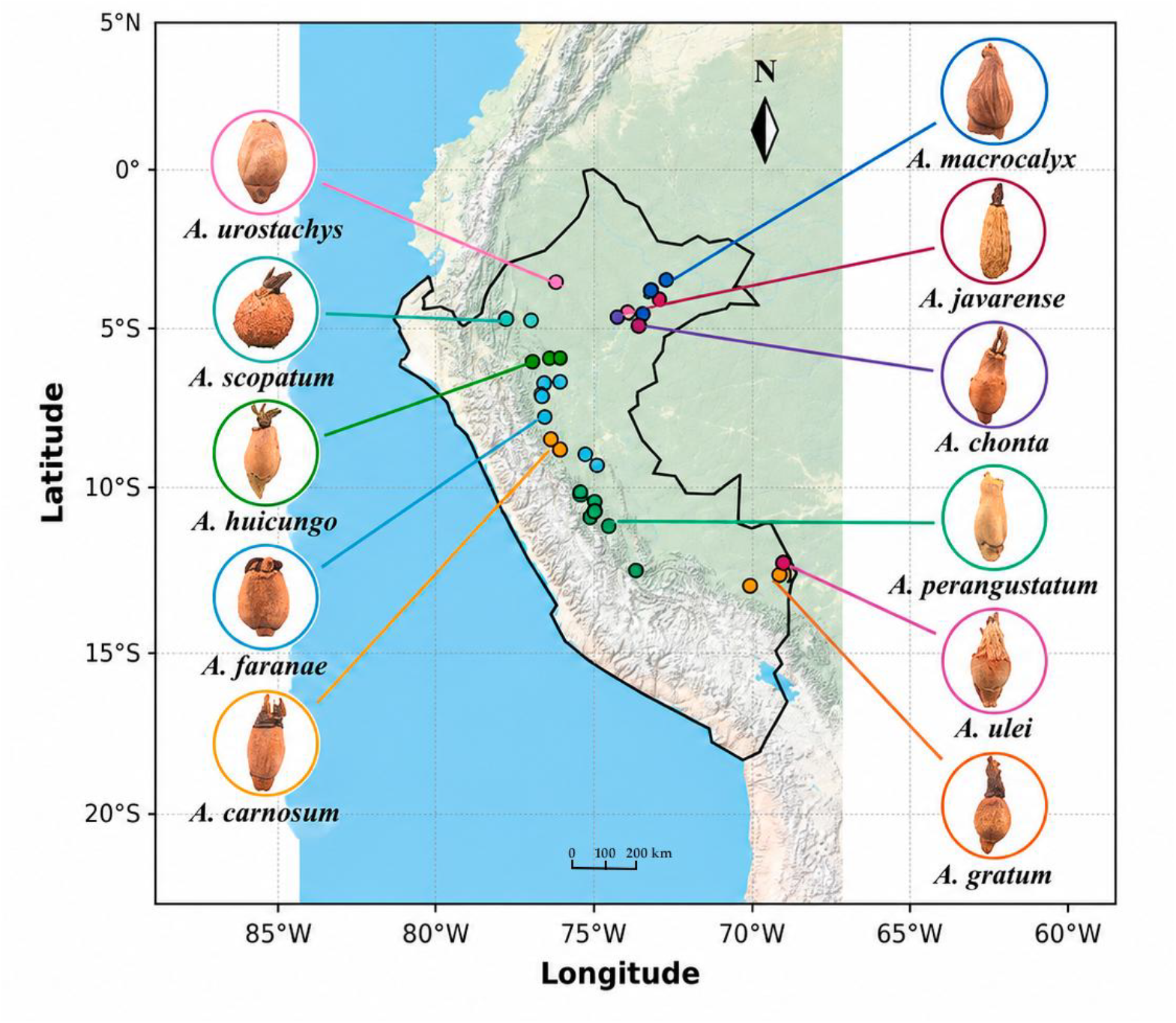
Geographic locations of the studied specimens of *Astrocaryum* sect. *Huicungo* in the Peruvian Amazon.

To guarantee the architectural representativeness of the inflorescences and avoid morphological alterations associated with maturation, individual selection was governed by two strict criteria: (i) possessing complete inflorescences that allowed for the equitable extraction of flowers from the basal, middle, and apical regions of the rachis; and (ii) verifying that the flowers had been collected during anthesis, given that fertilization induces allometric variations in perianth size. In total, 549 pistillate flowers sampled from 61 individuals were measured: *N* = 54 for *Astrocaryum carnosum*; *N* = 54 for *A. chonta*; *N* = 90 for *A. faranae*; *N* = 54 for *A. gratum*; *N* = 36 for *A. huicungo*; *N* = 27 for *A. javarense*; *N* = 54 for *A. macrocalyx*; *N* = 81 for *A. perangustatum*; *N* = 54 for *A. scopatum*; *N* = 27 for *A. ulei*; and *N* = 18 for *A. urostachys*.

### 4.2. Image Capture and Landmark Digitization

To expose the perianth contour and minimize visual artifacts, a superficial cleaning of the calyx and corolla was performed to remove prickles and bristles, thereby ensuring a clear visualization of the structure’s outline.

The floral structures were mounted on a support equipped with integrated lighting to eliminate shadows and were photographed alongside a metric scale. The anatomical orientation of each sample was standardized, strictly adhering to the original arrangement of the pedicel relative to the rachis.

Photographic captures were performed using a UNITRON Model 13435 trinocular zoom stereo microscope coupled with CaptaVision+ imaging software. To eliminate measurement bias, all images were recorded by a single operator [34]. To represent perianth geometry, a 2D configuration of 20 points (4 landmarks and 16 semilandmarks) was designed. The curvature was modeled utilizing 4 landmarks (points 1, 10, 11, and 20) and 16 semilandmarks (points 2 to 9 and 12 to 19) (Figure 7) [35]. Image file compilation into TPS format and subsequent manual digitization were executed using tpsUtil v.1.81 and tpsDig2 v.2.32 software, respectively [36].

**Figure 7.**
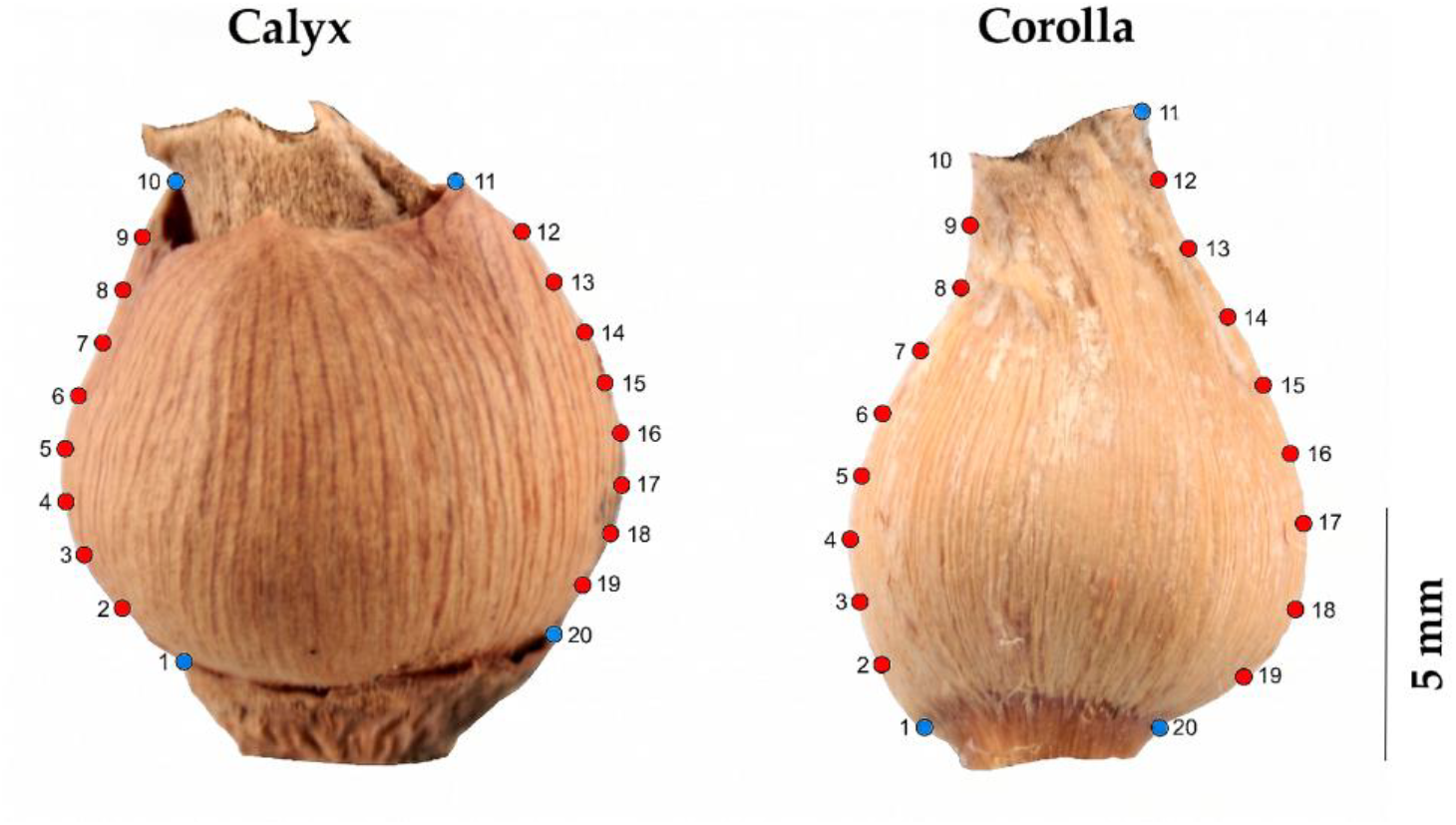
Configuration of landmarks (light blue) and semilandmarks (red) used to represent the shape of the calyx and corolla of the pistillate flower.

### 4.3. Data Analysis

Spatial processing and statistical analyses were conducted in R version 4.4.3 [37], implementing the specialized packages *geomorph* version 4.0.10 [9,38] and *Morpho* version 2.12 [39]. Initially, raw coordinate configurations were subjected to a Generalized Procrustes Analysis (GPA) to mathematically remove the effects of scale, position, and orientation [40,41]. To ensure statistical equivalence across the configuration, semilandmarks were mathematically slid along the contour curves applying the minimum bending energy criterion [11,42].

Because *Astrocaryum* pistillate flowers exhibit bilateral object symmetry in frontal view, shape variation was analytically partitioned [43]. Mirrored copies of the original coordinates were generated, and paired landmarks were relabeled to perform a joint Procrustes superimposition. This procedure allowed for the isolation of fluctuating and directional asymmetry from the symmetric component [16]. This symmetric component was used exclusively for all subsequent inter- and intraspecific analyses. Morphological trends were explored using a Principal Component Analysis (PCA), visualizing the trajectories of geometric change through thin-plate spline (TPS) deformation grids [40]. Morphological dispersion in the morphospace was evaluated by calculating the mean shapes for each species and generating PCA scatter plots with confidence ellipses. To quantify taxonomic differentiation, a Canonical Variate Analysis (CVA) was performed, and the robustness of the recovered morphological groups was assessed by extracting Mahalanobis distances subjected to an iterative permutation test (10,000 iterations) with a Bonferroni correction for multiple comparisons [13,44]. Furthermore, cross-validated classification rates (CCR) were computed to establish the assignment accuracy for each species. Additionally, the Mahalanobis distance matrices were used to construct phenetic similarity dendrograms using the UPGMA clustering algorithm. The entire statistical workflow was applied independently to the calyx and the corolla. Finally, a concatenated dataset (calyx + corolla) was analyzed to evaluate whether combining both floral structures improved taxonomic discrimination within section *Huicungo*.

## 5. Conclusions

Geometric morphometric analysis of the perianth in *Astrocaryum* sect. *Huicungo* pistillate flowers demonstrates that floral shape exhibits continuous variation and broad interspecific phenotypic overlap. Despite this high variability, the geometric architecture of the calyx and corolla retains a significant diagnostic signal that enables the statistical discrimination of most studied taxa.

Phenetic similarity relationships derived from perianth geometry are incongruent with traditional taxonomic subsections, supporting recent molecular evidence that challenges their monophyly. Although the isolated calyx conformation reveals a high geometric similarity among *Astrocaryum carnosum, A. faranae*, and *A. huicungo*, the combined perianth analysis successfully discriminates *A. carnosum* from the other two species. This intricate morphological continuum suggests limited character differentiation or very recent divergence rather than extensive evolutionary convergence, highlighting that some specific boundaries within the current classification remain diffuse.

Finally, these findings underscore that taxonomic delimitation in *Astrocaryum* should not rely exclusively on qualitative shape descriptors. Instead, it requires the integration of geometric morphometrics, ecological attributes, and genomic data within a comprehensive integrative framework to definitively resolve the systematics of this complex Amazonian group.

## Author Contributions

Conceptualization, B.M.; Methodology, G.M. and V.M.; formal analysis, G.M. and V.M.; investigation, G.M., V.M. and M.D.; data curation, G.M., V.M., M.D. and B.M.; visualization, G.M. and V.M.; writing—original draft preparation, G.M. and V.M.; writing—review and editing, B.M.; supervision, B.M. All authors have read and agreed to the published version of the manuscript.

## Data Availability Statement

The TPS landmark datasets and the R analysis workflow presented in this study are openly available in the Zenodo repository at https://doi.org/10.5281/zenodo.21176286.

## Acknowledgments

The authors express their sincere gratitude to the San Marcos Herbarium (USM) and the Department of Gymnosperms and Monocotyledons at the Museo de Historia Natural (MHN-UNMSM) for providing the facilities, infrastructure, and botanical specimens necessary for this study, as most of the research activities were fully developed within these departments.

